# Bmp8a Is a Novel Player in Regulation of Antiviral Immunity

**DOI:** 10.1101/2020.08.09.243519

**Authors:** Shenjie Zhong, Haoyi Li, Yun-Sheng Wang, Ying Wang, Guangdong Ji, Hong-Yan Li, Shicui Zhang, Zhenhui Liu

**Affiliations:** College of Marine Life Science and Institute of Evolution & Marine Biodiversity, Ocean University of China, Qingdao 266003, China; Laboratory for Marine Biology and Biotechnology, Pilot National Laboratory for Marine Science and Technology (Qingdao), Qingdao 266003, China

## Abstract

Bone morphogenetic protein (BMP) is a kind of classical multi-functional growth factor that plays a vital role in the formation and maintenance of bone, cartilage, muscle, blood vessels, and the regulation of energy balance. Whether BMP plays a role in antiviral immunity is unknown. Here we demonstrate that Bmp8a is a newly-identified positive regulator for antiviral immune responses. The *bmp8a*^−/−^ zebrafish, when infected with the viruses of GCRV, SVCV or TSVDV, show significantly reduced antiviral immunity, increased viral load and morbidity. We also show for the first time that Bmp8a interacts with Alk6a, which promotes the phosphorylation of Tbk1 and Irf3 through p38 MAPK pathway, and induces the production of type I IFNs in response to virus infection. Upon virus infection, *bmp8a* expression is activated through the binding of Stat1a/Stat1b to the GAS motifs in *bmp8a* promoter region, enlarging the antiviral innate immune signal. Our study uncovers a previously unrecognized role of Bmp8a in regulation of antiviral immune responses and provides a new target for controlling viral infection.

## Introduction

Viruses infect all groups of living things and produce a variety of diseases. Host cell detection of viral infection depends on the recognition of pathogen-associated molecular patterns (PAMPs) from viral proteins or nucleic acids by the pattern recognition receptors (PRRs) such as Toll-like receptors (TLRs), RIG-I-like receptors (RLRs), NOD-like receptors (NLRs) and cytoplasmic DNA or RNA sensors^1–6^. The PRRs, once stimulated by their appropriate ligands, trigger distinct intracellular signaling pathways that converge on the activation of IFN regulatory factor 3 (IRF3) and/or IFN regulatory factor 7 (IRF7), which then translocate into the nucleus and activate the production of interferons (IFNs) that play important roles in inhibiting virus replication^7,8^.

IFNs are classified into type I IFNs that include IFNα and IFNβ; type II IFN which has only a single member IFN-γ; and type III IFNs that consist of IFN-λs^9^. All these three types of IFNs trigger intracellular signaling cascades generally via the Janus kinase signal transducer and activator of transcription (JAK-STAT) pathway^10^. Canonically, type I IFNs initiate the signaling via binding to a heterodimeric receptor complex IFN-α/β receptor (IFNAR), inducing phosphorylation of transcription factors STAT1 and STAT2. The phosphorylated STATs form a trimolecular complex with IRF9, which translocates to the nucleus and binds to IFN-stimulated response elements (ISREs) of IFN-stimulated genes (ISGs), resulting in transcriptional activation of the target ISGs, including those encoding numerous cytokines and antiviral proteins^11–16^. In contrast, type II IFN binds to its unique receptor, IFN-γ receptor (IFNGR), resulting in the phosphorylation of STAT1 and the formation of STAT1 homodimers that recognize gamma-activated sequences (GASs) present in the promoter regions of IFN-γ-regulated genes^9^.

Bone morphogenetic proteins (BMPs) are potent growth factors belonging to the transforming growth factor beta (TGFβ) superfamily, which are intercellular signaling molecules with multiple functions in development and differentiation, such as skeletal formation and hematopoiesis, as well as lipid oxidation^17–21^. However, accumulating data suggests that BMPs also play important roles in the regulation of immune responses^22–27^. For example, BMP7 has been shown to drive Langerhans cell differentiation^28^. Interestingly, BMP signaling appears to have different responses in T cells: BMP4 and BMP6 were shown to promote T cell proliferation, whereas BMP2 was seen to inhibit T cell proliferation^29,30^. In addition, although BMPs were found to promote tumor growth by inhibiting the functions of cytotoxic T lymphocytes (CTLs) and dendritic cells (DCs), they were also shown to inhibit cancer cell proliferation and promote the activity of NK cells that typically display antitumour activity^31^. Therefore, it remains controversial regarding the functions of BMPs in immune responses.

In humans and mice, two closely related BMP8 genes, *BMP8A* and *BMP8B*, have been identified^32,33^. BMP8B has been found to play roles in the regulation of thermogenesis in mature brown adipose tissue and spermatogenesis in male germ cells, while BMP8A shown to closely correlate with the progression of spermatogenesis^34,35^. In contrast, only one *bmp8* gene, *bmp8a*, has been isolated in zebrafish, and shown to be involved in the regulation of lipid metabolism^36^. To date, whether Bmp8a plays a role in immune responses is completely unknown. The present study was thus performed to answer this question. We demonstrated that Bmp8a was a previously unrecognized regulator functioning in antiviral immune responses of zebrafish *Danio rerio*, providing a new target for control of viral infection.

## Results

### Bmp8a inhibited RNA virus replication *in vitro*

To explore the function of zebrafish Bmp8a in the antiviral immune response, we infected both the wild-type zebrafish liver (ZFL) cells and ZFL cells over-expressing bmp8a as well as the epithelioma papulosum cyprini (EPC) cells and EPC cells over-expressing bmp8a with Grass carp reovirus (GCRV), a dsRNA virus, and then monitored the cytopathic effect(CPE)and the viral titers of the supernatants. In response to GCRV challenge, an apparent CPE was observed in the control cells, while the bmp8a-overexpressing ZFL cells had much less CPE (Fig. 1a). In accordance with the CPE results, a significant decrease of the viral titers was observed in the supernatants of bmp8a-overexpressing ZFL cells (Fig. 1b). Similarly, the GCRV-induced CPE and GCRV yields of the supernatants were also markedly reduced in the bmp8a-overexpressing EPC cells (Fig. 1c, d). These suggested that Bmp8a suppressed the replication of the RNA virus in the cells.

**Fig. 1.**
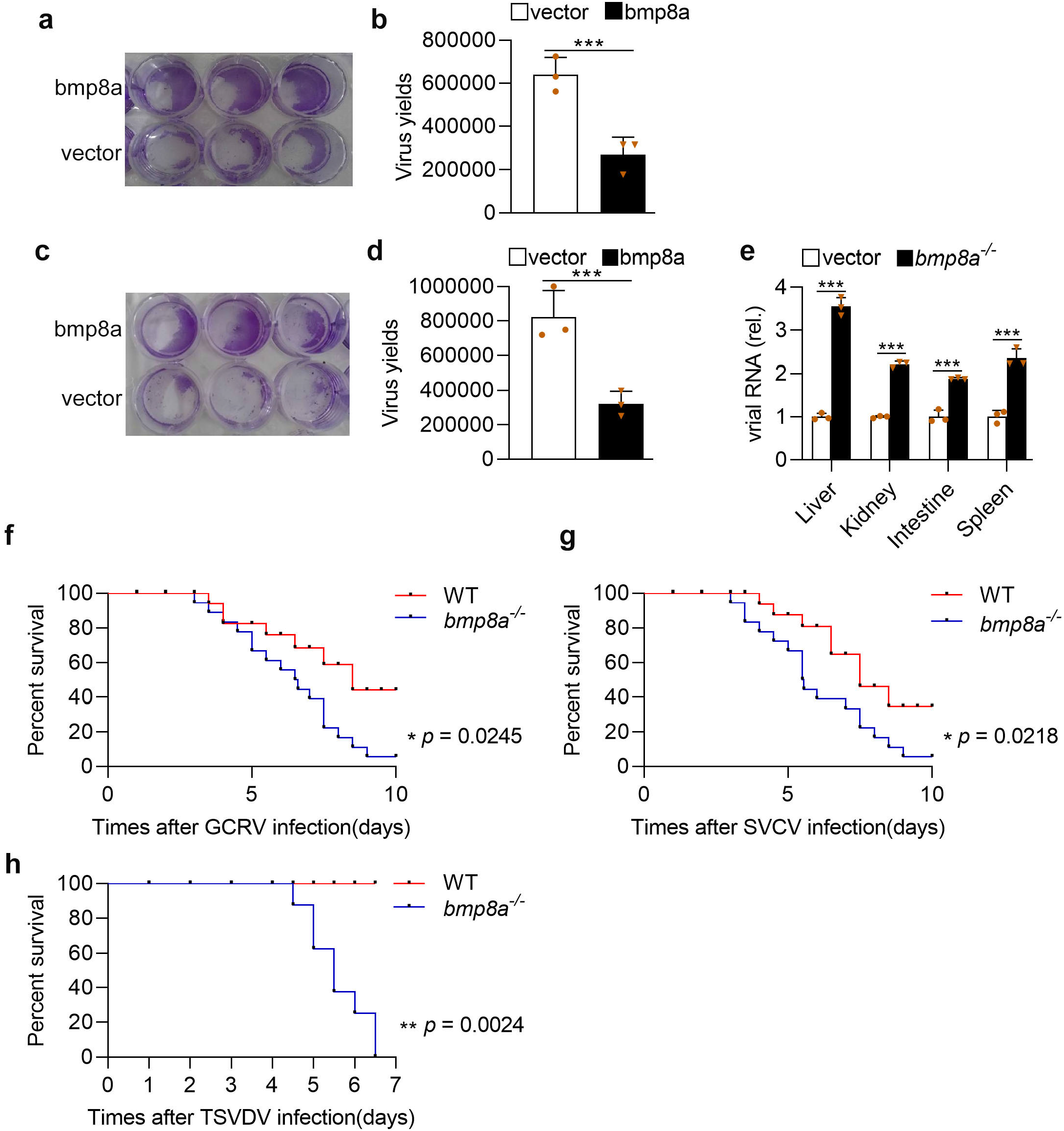
Bmp8a inhibits RNA virus replication both *in vitro* and *in vivo*. **a**, **b** ZFL cells were transfected with bmp8a (1μg) or empty vector (1μg), respectively. The cells were infected with GCRV (5 × 10^4^TCID_50_ per ml) at 24 h post-transfection, and the culture supernatants were collected at 72 h post-infection. The cell monolayers were fixed with 4% paraformaldehyde for 1 h and stained with 0.5% crystal violet for 2 h (**a**), and the viral titers of the supernatants were determined by TCID_50_ assays (**b**). **c**, **d** Similar as (**a**, **b**) but in EPC cells. **e** The expression of GCRV RNA in the liver, kidney, intestine, and spleen from wild-type (WT) or *bmp8a*^−/−^ zebrafish injected i.p. with 50 μl of GCRV (10^8^TCID_50_ per ml) for 72 h. **f**, **g** Kaplan–Meier analysis of the overall survival of WT (*n* = 20) or *bmp8a*^−/−^ zebrafish (*n* = 20) which were injected i.p. with 50 μl of GCRV (10^8^TCID_50_ per ml) or SVCV (10^8^TCID_50_ per ml) and monitored every 12 h after infection. **h** Kaplan– Meier analysis of the overall survival of WT (*n* = 8) or *bmp8a*^−/−^ zebrafish (*n* = 8) which were injected i.p. with 50 μl of TSVDV (crude virus extracts filtered by a 0.45 μm microporous membrane) and monitored every 12 h after infection. The expression of zebrafish *actb1* was used as an internal control for the qRT-PCR. Data were from three independent experiments (**a–e**) or two independent experiments (**f-h**). Data were analyzed by Student’s *t*-test (two-tailed) or log-rank (Mantel–Cox) test and were presented as mean ± SD (\**p* < 0.05, \*\*\**p* < 0.001). Source data are provided as a Source Data file.

### Bmp8a was involved in antiviral responses *in vivo*

To test the function of Bmp8a in host defense against virus infection *in vivo*, we generated *bmp8a* deficient (*bmp8a*^−/−^) zebrafish using the TALEN approach (Supplementary Fig. 1a), which caused 7 nucleotides deletion in the exon 4 (Supplementary Fig. 1b, c). We challenged both the wild-type and *bmp8a*^−/−^ mutant zebrafish by intraperitoneal injection with GCRV. Compared with wild-type zebrafish, the levels of GCRV RNA in the liver, kidney, intestine, and spleen of *bmp8a*^−/−^ zebrafish were considerably increased (Fig. 1e). Accordingly, *bmp8a*^−/−^ zebrafish exhibited significantly reduced survival rate than wild-type zebrafish upon GCRV infection (Fig. 1f). When the wild-type and *bmp8a*^−/−^ zebrafish were similarly challenged with spring viremia of carp virus (SVCV), a negative ssRNA virus, the survival rate of *bmp8a*^−/−^ zebrafish was also remarkably lower than that of wild-type zebrafish (Fig. 1g). Moreover, when the wild-type and *bmp8a*^−/−^ zebrafish were infected with turbot skin verruca disease virus (TSVDV) isolated from skin (Supplementary Fig. 2), mortality occurred in *bmp8a*^−/−^ zebrafish at 108 h postinfection, and all the fish died at 156 h post infection, while none of the wild type zebrafish were dead during the period (Fig. 1h). These data indicated that *bmp8a*^−/−^ zebrafish were more susceptible to virus infection than wild-type zebrafish, suggesting involvement of Bmp8a in antiviral immune responses *in vivo*.

### Bmp8a promoted expression of antiviral genes

In zebrafish, type I IFNs contained 4 members *ifnφ1*, *ifnφ2*, *ifnφ3*, and *ifnφ4*, among which *ifnφ1* and *ifnφ3* were shown to be involved in the antiviral response. In contrast, in carp, only one type I IFN was identified to fulfill antiviral role in EPC cell^37,38^. To pinpoint the role of Bmp8a in the antiviral immune response, both the wild-type and *bmp8a*^−/−^ zebrafish were injected intraperitoneally with GCRV, and the expression of type I IFN genes and the antiviral protein gene *mx* were detected using quantitative reverse transcription PCR (qRT-PCR). We found that the expression of both *ifnφ1* and *ifnφ3* as well as *mx* was all significantly down-regulated in the liver, kidney, intestine, and spleen of *bmp8a*^−/−^ zebrafish, compared with those of wild-type fish (Fig. 2a-d). We then over-expressed bmp8a in ZFL cells and found that ZFL cells with over-expressed Bmp8a showed markedly higher expression of *ifnφ1*, *ifnφ3*, and *mx* than that of control cells, infected with or without GCRV (Fig. 2e-h). In addition, *bmp8a* knockdown caused remarkably decreased expression of *ifnφ1*, *ifnφ3*, and *mx* in ZFL cells (Fig. 2i-l). Similarly, the expression of *ifn* and *mx* in EPC cells with over-expressed Bmp8a was also remarkably up-regulated than that in control cells (Fig. 2m-p). Moreover, EPC cells cotransfected with bmp8a expressing plasmid and IFNφ1, IFNφ3, or EPC IFN promoter-driven luciferase plasmid, followed by infection with or without GCRV, had markedly increased intracellular IFNφ1, IFNφ3, or EPC IFN promoter-driven luciferase activities (Fig. 2q-s). All these data denoted that Bmp8a promoted the expression of both type I IFN and *mx* genes.

**Fig. 2.**
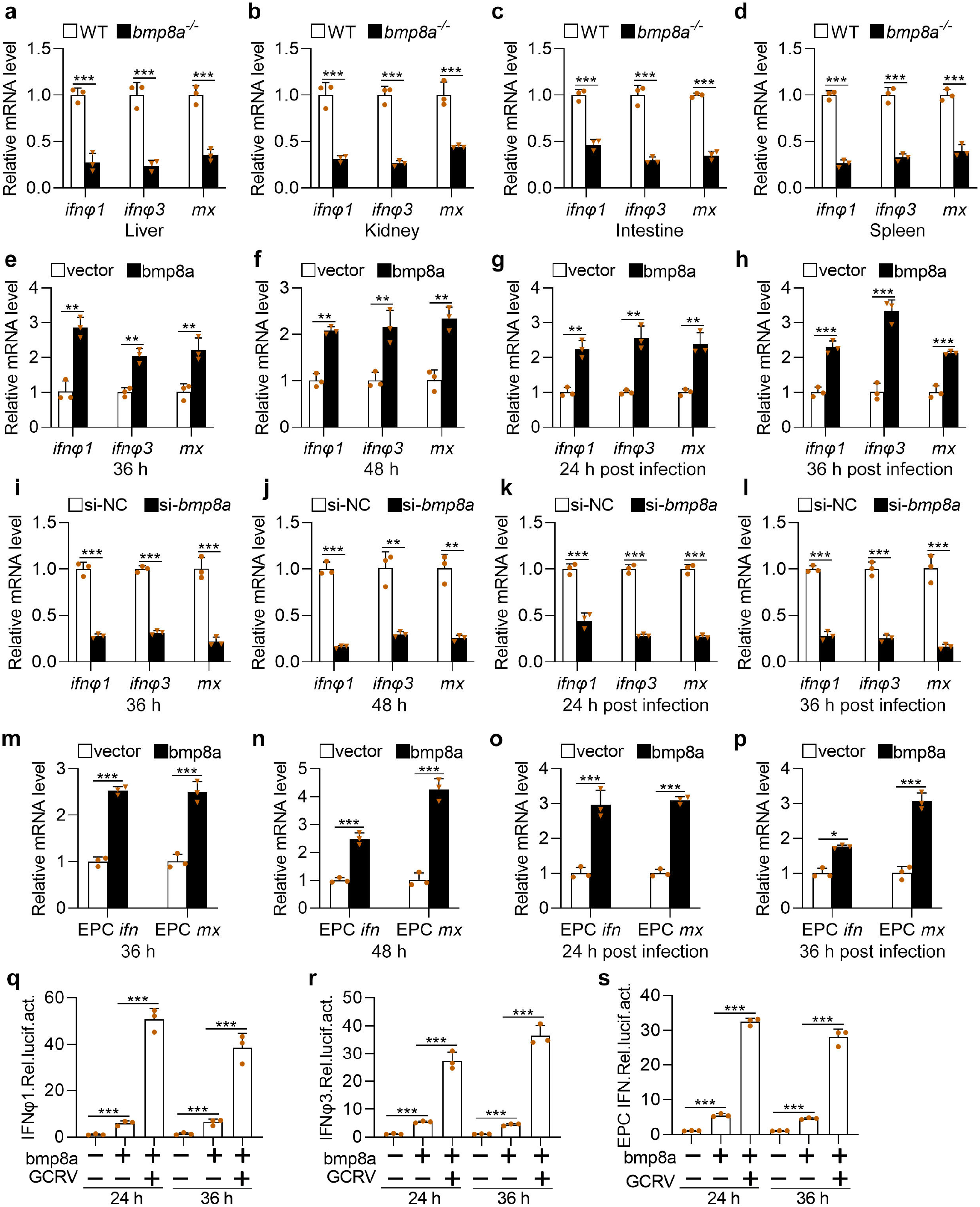
Bmp8a promotes antiviral innate immune responses. **a-d** Expression of *ifnφ1*, *ifnφ3*, and *mx* mRNA in the liver, kidney, intestine, and spleen from WT or *bmp8a*^−/−^ zebrafish injected i.p. with 50 μl of GCRV (10^8^TCID_50_ per ml) for 72 h. **e**, **f** Expression of *ifnφ1*, *ifnφ3*, and *mx* mRNA after transfected with bmp8a (2 μg) or empty vector (2 μg) in ZFL cells. The cells were collected at 36 h (**e**) or 48 h (**f**) post transfection. **g, h** Expression of *ifnφ1*, *ifnφ3*, and *mx* mRNA after transfected with bmp8a (2 μg) or empty vector (2 μg) in ZFL cells for 24 h, followed by infection with GCRV for another 24 h (**g**) or 36 h (**h**). **i**, **l** Expression of *ifnφ1*, *ifnφ3*, and *mx* mRNA after *bmp8a* knockdown in ZFL cells. The cells were collected at 36 h (**i**) or 48 h (**l**) post knockdown. **k**, **l** Expression of *ifnφ1*, *ifnφ3*, and *mx* mRNA after knockdown *bmp8a* in ZFL cells for 24 h, followed by infection with GCRV for another 24h (**k**) or 36 h (**l**). **m**, **n** Expression of EPC *ifn* and EPC *mx* mRNA after transfected with bmp8a (2 μg) or empty vector (2 μg) in EPC cells. The cells were collected at 36 h (**m**) or 48 h (**n**) post transfection. **o**, **p** Expression of EPC *ifn* and EPC *mx* mRNA after transfected with bmp8a (2 μg) or empty vector (2 μg) in EPC cells for 24 h, followed by infection with GCRV for another 24 h (**o**) or 36 h (**p**). The expression of zebrafish *actb1* or EPC *actin* was used as an internal control for the qRT-PCR. **q-s** EPC cells were transfected with IFNφ1pro-luc (200 ng, **q**), IFNφ3pro-luc (200 ng, **r**) or EPC IFN pro-luc (200 ng, **s**) respectively, with or without bmp8a (200 ng), followed by infection with GCRV. At the indicated time points, cells were collected for luciferase assays. *Renilla* luciferase was used as the internal control. Data were from three independent experiments and were analyzed by Student’s *t*-test (two-tailed) and were presented as mean ± SD (\*\**p* < 0.01, \*\*\**p* < 0.001). Source data are provided as a Source Data file.

### Bmp8a activated Tbk1-Irf3-Ifn antiviral signaling via p38 MAPK pathway

To putatively examine the molecular mechanism by which Bmp8a stimulates *ifn* and *mx* expression, we carried out transcriptome analysis of the livers of *bmp8a*^−/−^ mutant and wild-type zebrafish, that both had been intraperitoneally injected with GCRV. Kyoto Encyclopedia of Genes and Genomes (KEGG) analyses revealed that, compared with wild-type zebrafish, the down-regulated genes in the mutant were remarkably enriched in the RIG-I-like receptor signaling pathway, NF-kappa B signaling pathway, TNF signaling pathway, and protein digestion and absorption that function in the antiviral immune process (Supplementary Fig. 3). In agreement with transcriptome analysis, we found that the expressions of *tbk1*, *irf3*, and *irf7* were significantly upregulated in bmp8a-overexpressing ZFL cells than control cells, infected with or without GCRV (Fig. 3a-d). Similar expression patterns of *tbk1*, *irf3*, and *irf7* were also observed in EPC cells, infected with or without GCRV (Fig. 3e-h). In addition, *bmp8a* knockdown resulted in considerably reduced expression of *tbk1*, *irf3*, and *irf7* in ZFL cells than control cells, infected with or without GCRV (Fig. 3i-l). We then evaluated the activation of Tbk1 and Irf3 by immunoblot assays in both bmp8a-overexpressing and wild-type ZFL cells as well as EPC cells, infected with or without GCRV. It was found that phosphorylation levels of Tbk1 and Irf3 were significantly increased in both types of the cells with overexpressed Bmp8a (Fig. 3m-p). Moreover, *bmp8a* knockdown markedly reduced the phosphorylation levels of Tbk1 and Irf3 in ZFL cells, infected with or without GCRV (Fig. 3q, r). Furthermore, in luciferase activity assays, transfection of dominant negative mutations of *tbk1* (tbk1-K38M), *irf3* (irf3DN), or *irf7* (irf7DN) in EPC cells induced a significant loss in the ability of Bmp8a to activate the IFN promoter (Fig. 3s-u). This was further supported by the observations *in vivo* that the expression of *tbk1*, *irf3*, and *irf7* in the liver, kidney, intestine, and spleen was significantly downregulated in *bmp8a*^−/−^ mutant than that in wild-type zebrafish (Fig. 3v-y). These data together revealed that Bmp8a induced *ifn* and *mx* expression via promoting Tbk1-Irf3-Ifn antiviral signaling.

**Fig. 3.**
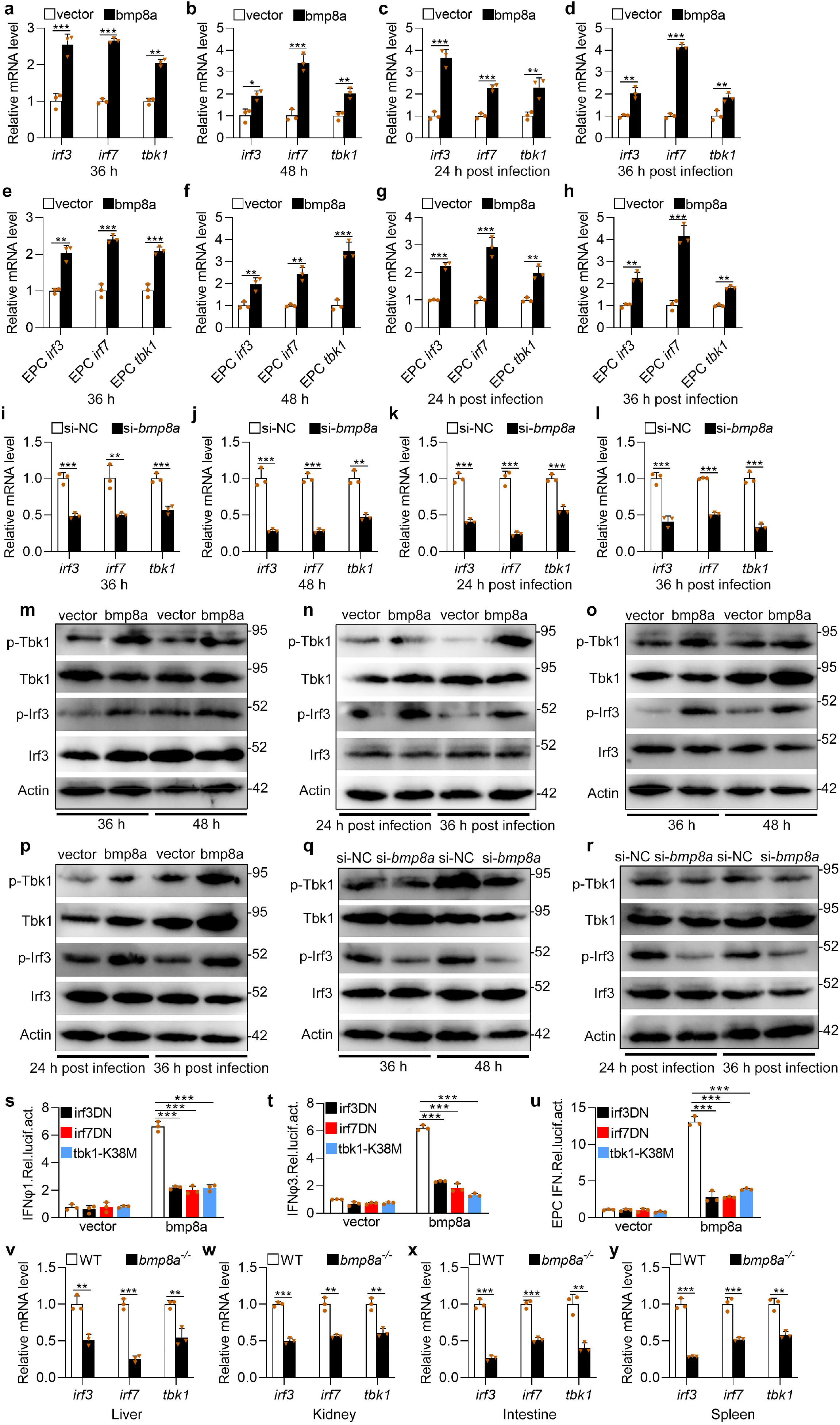
Bmp8a increases Tbk1-Irf3-Ifn antiviral signaling. **a**, **b**, **e**, **f** Expression of *irf3*, *irf7*, and *tbk1* mRNA after transfected with 2 μg bmp8a or empty vector in ZFL (**a**, **b)** or EPC (**e**, **f)** cells. The cells were collected at 36 h (**a**, **e**) or 48 h (**b**, **f**) post transfection. **c**, **d**, **g**, **h** Expression of *irf3*, *irf7*, and *tbk1* mRNA after transfected with 2 μg bmp8a or empty vector in ZFL (**c**, **d)** or EPC **(g**, **h)**cells for 24 h, followed by infection with GCRV for another 24 h (**c**, **g**) or 36 h (**d**, **h**). **i, j, k, l** Expression of *irf3*, *irf7*, and *tbk1* mRNA after *bmp8a* knockdown in ZFL cells. The cells were collected at 36 h (**i**) and 48 h (**j**) post knockdown or at 24 h (**k**) and 36 h (**l**) post infected with GCRV. **m**, **o** Immunoblot analysis of phosphorylated (p-) Tbk1 and Irf3 after transfected with 2 μg bmp8a or empty vector in ZFL (**m**) or EPC (**o**) cells. The cells were collected at 36 h or 48 h post transfection for Immunoblot analysis. **n**, **p** Immunoblot analysis of phosphorylated (p-) Tbk1 and Irf3 after transfected with 2 μg bmp8a or empty vector in ZFL (**n**) or EPC (**p**) cells for 24 h, followed by infection with GCRV for another 24 h or 36 h. **q**, **r** Immunoblot analysis of phosphorylated (p-) TBK1 and IRF3 after *bmp8a* knockdown in ZFL cells. The cells were collected at 36 h and 48 h post knockdown or at 24 h and 36 h post infected with GCRV. **s-u** EPC cells were cotransfected with IFN-φ1pro-luc (200 ng, **s**), IFN-φ3pro-luc (200 ng, **t**) or EPC IFNpro-luc (200ng, **u**), and bmp8a (100 ng) together with each of the dominant negative plasmids including tbk1-K38M (100 ng), irf3DN (100 ng) and irf7DN (100 ng). At 48 h post transfection, the cells were collected for luciferase assays. *Renilla* luciferase was used as the internal control. **v-y** Expression of *irf3*, *irf7*, and *tbk1* mRNA in the liver, kidney, intestine, and spleen from WT or *bmp8a*^−/−^ zebrafish injected i.p. with 50 μl of GCRV (10^8^TCID_50_ per ml). The expression of zebrafish *actb1* or EPC *actin* was used as an internal control for the qRT-PCR. Data were from three independent experiments and were analyzed by Student’s *t*-test (two-tailed) and were presented as mean ± SD (\*\**p* < 0.01, \*\*\**p* < 0.001). Source data are provided as a Source Data file.

BMPs transmit signals through smad1/5/8, smad2/3, ERK, JNK, or p38 MAPK pathways^39–41^. Thus, the effects of p38 MAPK inhibitor SB203580, JNK inhibitor SP600125, MEK1/2 inhibitor U0126, SMAD1/5/8 inhibitor DMH1, and smad2/3 inhibitor TP0427736 HCl on the expression of *ifn* was tested in bmp8a-overexpressing ZFL cells and EPC cells. We have shown above that Bmp8a overexpression increased the expression of *ifn* (Fig. 2). Here we found that only p38 MAPK inhibitor SB203580 significantly reduced the expression of *ifnφ1* and *ifnφ3* in ZFL cells and the expression of *ifn* in EPC cells (Fig. 4a-c). Furthermore, the IFN promoter-driven luciferase assays revealed that the luciferase activities were markedly reduced in EPC cells upon treatment with p38 MAPK inhibitor SB203580 (Fig. 4d-f), consistent with the observation that Bmp8a induced the expression of *ifn* through p38 MAPK pathway. Taken the previous report that the MAPK signaling increased the phosphorylation of TBK1 and IRF3 upon viral infection into consideration^42^, we suggested that Bmp8a activated Tbk1-Irf3-Ifn antiviral signaling via p38 MAPK pathway.

**Fig. 4.**
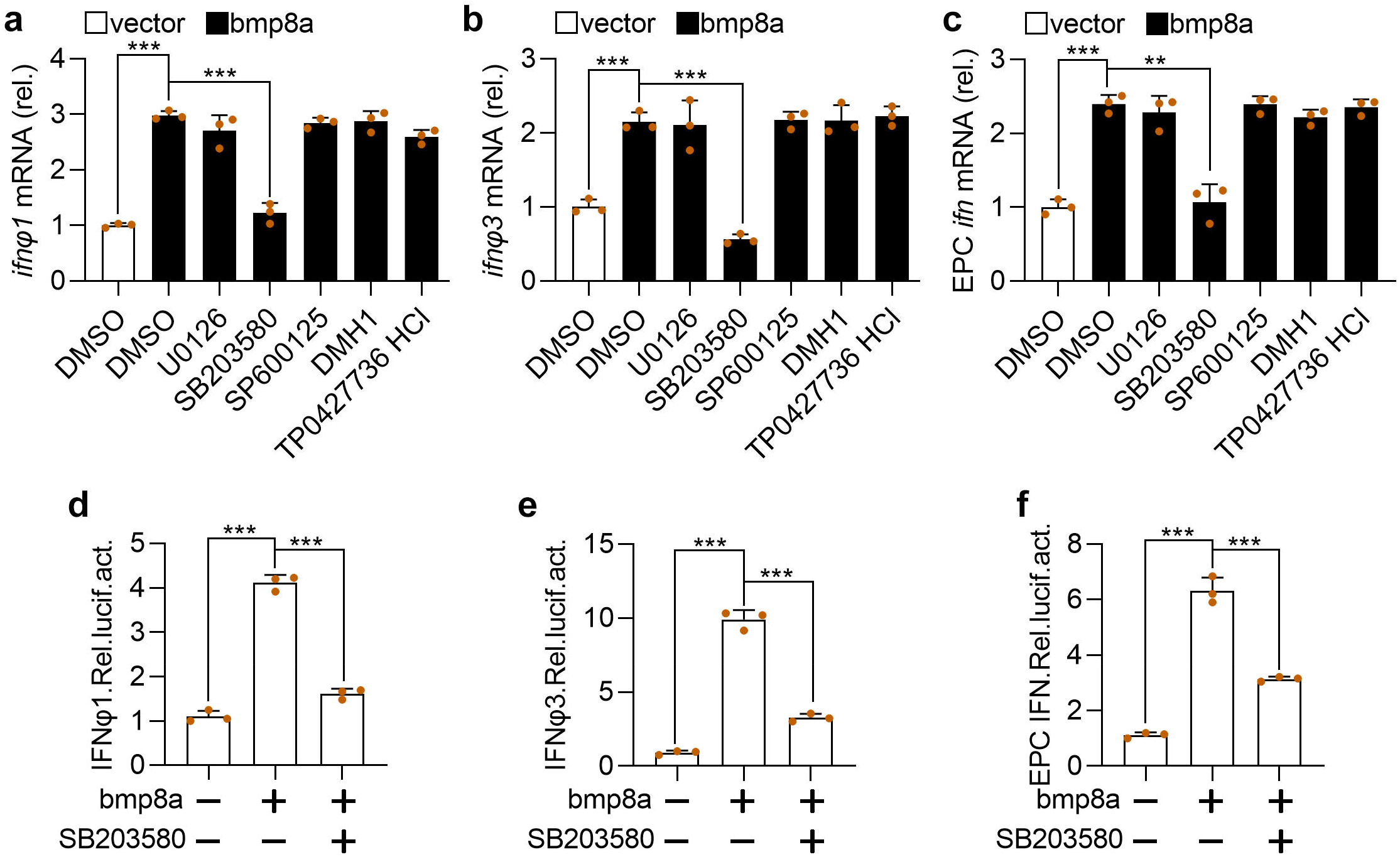
Bmp8a promotes the IFN expression via p38 MAPK pathway. **a**, **b** Expression of *ifnφ1* (**a**) and *ifnφ3* (**b**) mRNA after transfected with bmp8a (2 μg) in ZFL cells for 24 h, followed by treatment with SB203580, SP600125, U0126, DMH1 and TP0427736 HCl for another 24 h. **c** similar as (**a**, **b**) but in EPC cells. The expression of zebrafish *actb1* or EPC *actin* was used as an internal control for the qRT-PCR. **d-f** EPC cells were cotransfected with IFNφ1pro-luc (200 ng, **d**), IFNφ3pro-luc (200 ng, **e**) or EPC IFNpro-luc (200 ng, **f**), pRL-TK (20 ng) together with bmp8a (200 ng) or empty vector (200 ng), respectively. At 24 h post transfection, cells were treated with or without SB203580 for another 24 h and then harvested for detection of luciferase activity. *Renilla* luciferase was used as the internal control. Data were from three independent experiments and were analyzed by Student’s *t*-test (two-tailed) and were presented as mean ± SD (\*\**p* < 0.01, \*\*\**p* < 0.001). Source data are provided as a Source Data file.

### BMP type I receptor Alk6a interacted with Bmp8a and participated in antiviral immunity

BMPs exert their biological effects through the sequential activation of two types of transmembrane receptors, namely BMP receptor type I and BMP receptor type II, that both possess intrinsic serine/threonine kinase activity^43,44^. To detect which receptor is involved in the regulation of the antiviral immune responses, the expression of both type I receptor genes (*alk2*, *alk3*, *alk6a*) and type II receptor genes (*bmpr2a*, *bmpr2b*, *actr2a*, *actr2b*) in ZFL cells transfected with poly(I:C) or infected with GCRV were measured. Among these receptor genes, *alk6a* showed the highest expression upon poly(I:C) or virus challenge (Fig. 5a, b). We also overexpressed all these receptors in ZFL and EPC cells, and examined the expression of *ifn* in these cells. The upregulation of *ifnφ1* was only found in the alk6a-overexpressing ZFL cells (Fig. 5c). Moreover, the overexpression of *alk6a* or *alk2* resulted in significantly increased expression of *ifnφ3* in ZFL cells and *ifn* in EPC cells, with overexpressed *alk6a* being more effective (Fig. 5d, e). Based on these findings, the function of Alk6a was selected for further test.

**Fig. 5.**
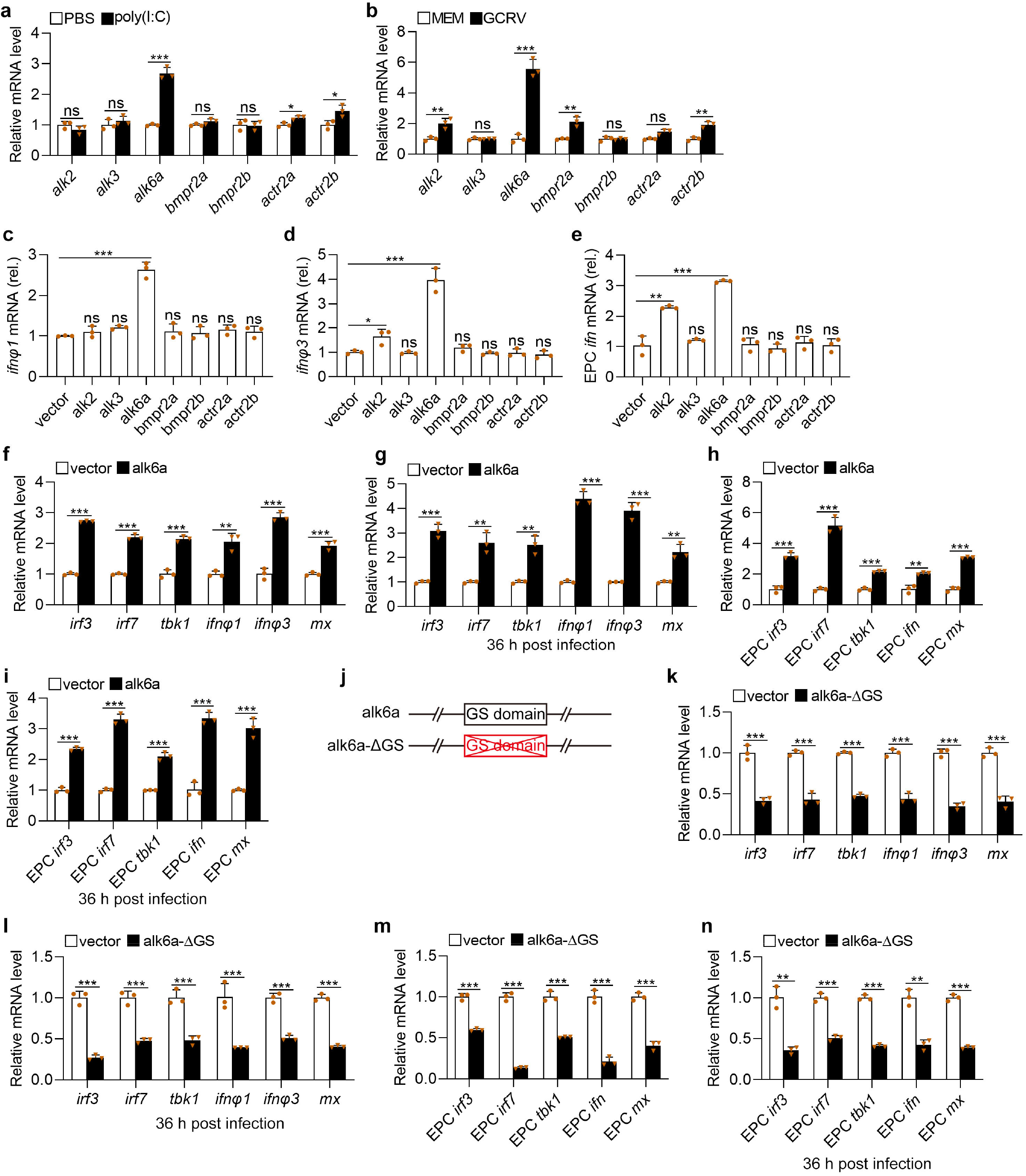
Alk6a is involved in the antiviral innate immune responses. **a**, **b** Expression of *alk2*, *alk3*, *alk6a*, *bmpr2a*, *bmpr2b*, *actr2a*, *actr2b* mRNA in ZFL cells stimulated with GCRV (5 × 10^4^TCID_50_ per ml, **a**) or poly(I:C) (2 μg/ml, **b**) for 48 h. **c-e** Expression of *ifnφ1* (**c**) and *ifnφ3* (**d**) mRNA in ZFL cells or EPC *ifn* (**e**) in EPC cells which were transfected with 2 μg of alk2, alk3, alk6a, bmpr2a, bmpr2b, actr2a, actr2b or empty vector for 48 h. **f, h** Expression of *irf3*, *irf7*, *tbk1*, *ifn* (or *ifnφ1* and *ifnφ3*), and *mx* mRNA after transfected with 2 μg of pcDNA3.1-alk6a or empty vector in ZFL (**f**) or EPC (**h**) cells for 48 h. **g, i** Expression of *irf3*, *irf7*, *tbk1*, *ifn* (or *ifnφ1* and *ifnφ3*), and *mx* mRNA after transfected with 2 μg of of pcDNA3.1-alk6a or empty vector in ZFL (**g**) or EPC (**i**) cells for 24 h, followed by infection with GCRV for another 36 h. **j** Schematic drawing of the alk6a-ΔGS mutation that the GS domain of Alk6a was deleted. **k, m** Expression of *irf3*, *irf7*, *tbk1*, *ifn* (or *ifnφ1* and *ifnφ3*), and *mx* mRNA after transfected with 2 μg pcDNA3.1-alk6a-ΔGS or empty vector in ZFL (**k**) or EPC (**m**) cells for 48 h. **l, n** Expression of *irf3*, *irf7*, *tbk1*, *ifn* (or *ifnφ1* and *ifnφ3*), and *mx* mRNA after transfected with 2 μg pcDNA3.1-alk6a-ΔGS or empty vector in ZFL (**l**) or EPC (**n**) cells for 24 h, followed by infection with GCRV for another 36 h. The expression of zebrafish *actb1* or EPC *actin* was used as an internal control for the qRT-PCR. Data were from three independent experiments and were analyzed by Student’s *t*-test (two-tailed) and were presented as mean ± SD (\**p* < 0.05, \*\**p* < 0.01, *** *p* < 0.001, ns means no significant difference). Source data are provided as a Source Data file.

Compared to wild-type ZFL cells, the expressions of antiviral protein genes *ifnφ1, ifnφ3* and *mx* as well as the antiviral signaling genes *tbk1*, *irf3*, and *irf7* were all markedly upregulated in alk6a-overexpressed ZFL cells with or without GCRV infection (Fig. 5f, g). The similar results were also observed in the alk6a-overexpressed EPC cells (Fig. 5h, i). Because the phosphorylation of Gly-Ser (GS) domain of the type I receptor is required for its activation, we thus constructed dominant-negative mutant of alk6a (alk6a-ΔGS) plasmid (Fig. 5j), and then transferred it into both ZFL and EPC cells. It revealed that the dominant-negative mutation of *alk6a* significantly reduced the expression of *mx, ifn* (*ifnφ1* and *ifnφ3* in ZFL cells), *tbk1*, *irf3*, and *irf7* in ZFL and EPC cells, with or without GCRV infection (Fig. 5k-n). The data suggested that Alk6a was apparently involved in the Tbk1-Irf3/7-Ifn antiviral signaling.

Considering we have revealed that Bmp8a activated Tbk1-Irf3-Ifn antiviral signaling via p38 MAPK pathway, we then examined whether the the levels of phosphorylated p38 MAPK protein were regulated by Alk6a. Clearly, compared to wild-type ZFL or EPC cells, phosphorylation levels of p38 MAPK were significantly increased in alk6a-overexpressed both types of the cells with or without GCRV infection (Fig. 6a-d). Moreover, dominant-negative mutation of *alk6a* significantly reduced the phosphorylation levels of p38 MAPK in ZFL or EPC cells, with or without GCRV infection (Fig. 6e-h). Thus, Alk6a was shown to be involved in the phosphorylation of p38 MAPK. Next, we investigated whether the antiviral immune responses of Bmp8a were mediated by Alk6a. In the IFN promoter-driven luciferase assays, it was found that the ability of Bmp8a to activate the IFN promoter was markedly blocked after the dominant-negative mutation of *alk6a* in EPC cells (Fig. 6i-k). Importantly, co-immunoprecipitation (Co-IP) experiments confirmed that Bmp8a interacted with Alk6a (Fig. 6l). Collectively, our data demonstrated that BMP type I receptor Alk6a interacted with Bmp8a, and participated in the antiviral immunity. To our best knowledge, Alk6a is the first BMP receptor identified thus far which is being directly required for antiviral immune responses.

**Fig. 6.**
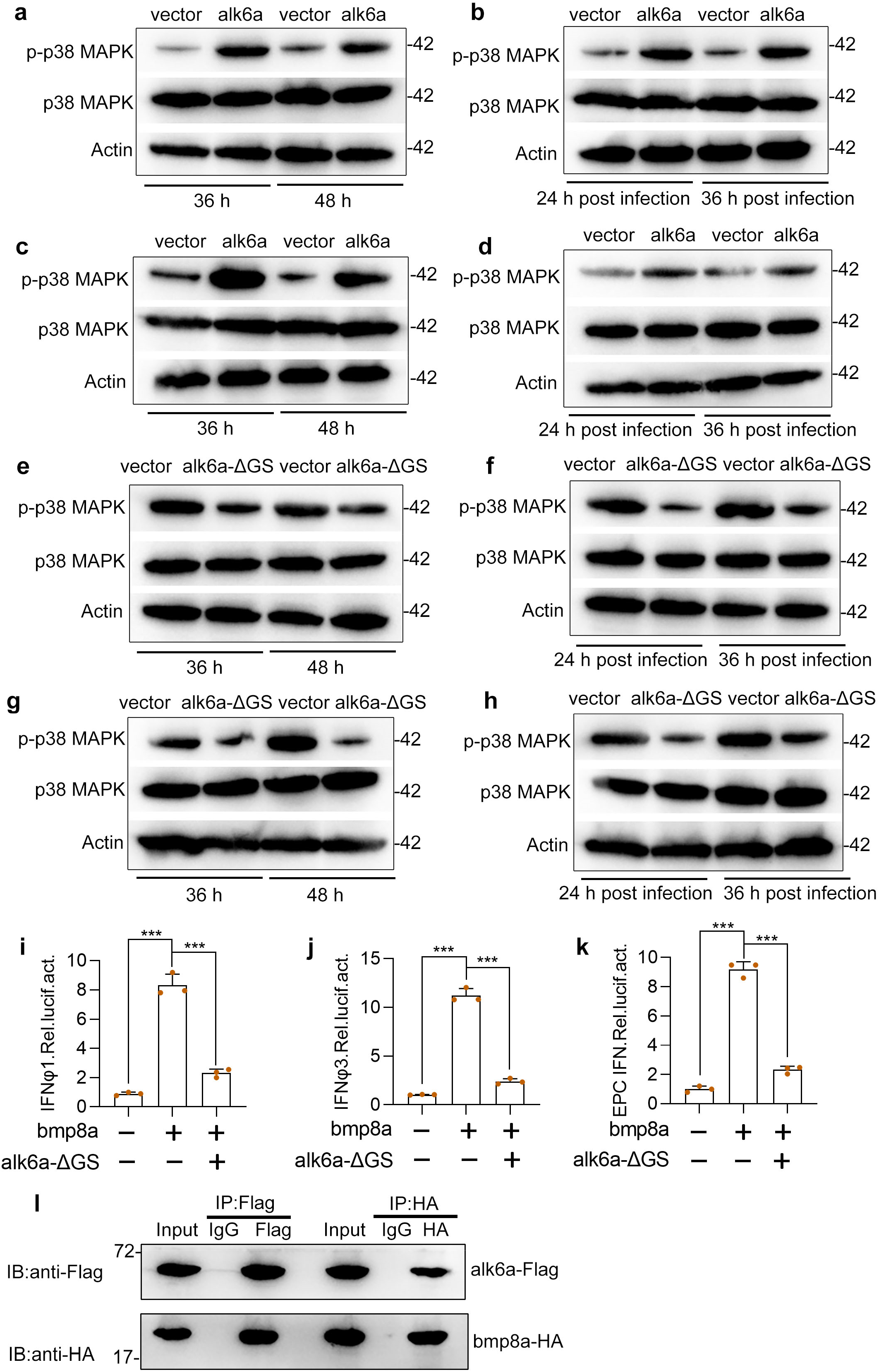
Bmp8a activates the IFN expression through Alk6a. **a**, **c** Immunoblot analysis of phosphorylated (p-) p38 MAPK after transfected with 2 μg alk6a or empty vector in ZFL (**a**) or EPC (**c**) cells. The cells were collected at 36 h or 48 h post transfection for Immunoblot analysis. **b**, **d** Immunoblot analysis of phosphorylated (p-) p38 MAPK after transfected with 2 μg alk6a or empty vector in ZFL (**b**) or EPC (**d**) cells for 24 h, followed by infection with GCRV (5 × 10^4^TCID_50_ per ml) for another 24 h or 36 h. **e, g** Immunoblot analysis of phosphorylated (p-) p38 MAPK after transfected with 2 μg alk6a-ΔGS or empty vector in ZFL (**e**) or EPC (**g**) cells. The cells were collected at 36 h or 48 h post transfection for Immunoblot analysis. **f**, **h** Immunoblot analysis of phosphorylated (p-) p38 MAPK after transfected with 2 μg alk6a or empty vector in ZFL (**f**) or EPC (**h**) cells for 24 h, followed by infection with GCRV (5 × 10^4^TCID_50_ per ml) for another 24 h or 36 h. **i-k** EPC cells were cotransfected with IFN-φ1pro-luc (200 ng, **i**), IFN-φ3pro-luc (200 ng, **j**) or EPC IFNpro-luc (200ng, **k**), and bmp8a (100 ng) together with or without the dominant negative plasmids alk6a-ΔGS (100 ng). At 48 h post transfection, the cells were collected for luciferase assays. *Renilla* luciferase was used as the internal control. **l** Co-immunoprecipitation and immunoblot analysis of EPC cells cotransfected with alk6a-Flag (1 μg) and bmp8a-HA (1μg). Data were from three independent experiments and were analyzed by Student’s *t*-test (two-tailed) and were presented as mean ± SD (*** *p* < 0.001). Source data are provided as a Source Data file.

### Virus induced expression of *bmp8a*

Next, we measured the expression of *bmp8a* in ZFL cells upon virus infection. GCRV significantly induced the expression of *bmp8a* at 12 h post-stimulation, and reached a peak at 48 h (Fig. 7a). We also detected the expression of *bmp8a* upon infection with GCRV or poly(I:C) *in vivo*. The expression of *bmp8a* in the liver, kidney, intestine, and spleen of zebrafish infected with GCRV or poly(I:C) was all significantly elevated (Fig. 7b-i). This indicated that the expression of *bmp8a* was inducible by infection with virus or its mimic poly(I:C).

**Fig. 7.**
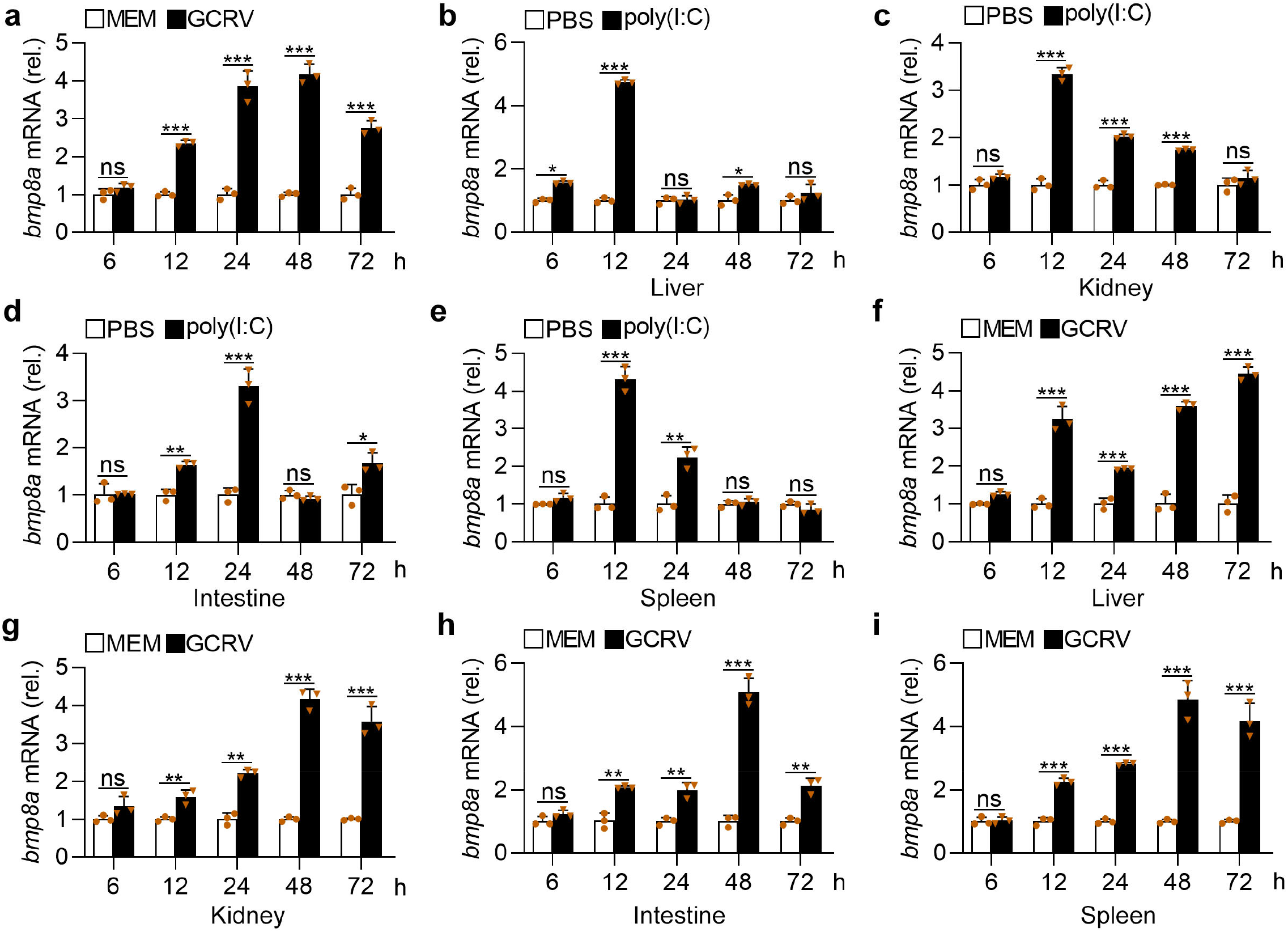
GCRV or poly(I:C) increases the expression of *bmp8a*. **a** Expression of *bmp8a* mRNA in ZFL cells after infected with GCRV (5 × 10^4^TCID_50_ per ml) for 48 h. **b-e** Expression of *bmp8a* mRNA in the liver (**b**), spleen (**c**), intestine (**d**), and kidney (**e**) from zebrafish injected i.p. with poly(I:C) (10 μg/fish). **f-i** Expression of *bmp8a* mRNA in the liver (**f**), spleen (**g**), intestine (**h**), and kidney (**i**) from zebrafish injected i.p. with 50μl of GCRV (10^8^TCID_50_ per ml). Zebrafish injected i.p. with PBS or MEM were used as the control. The expression of *actb1* served as a control for the qRT-PCR. Data were from three independent experiments and were analyzed by Student’s *t*-test (two-tailed) and were presented as mean ± SD (\**p* < 0.05, \*\**p* < 0.01, \*\*\**p* < 0.001, ns means no significant difference). Source data are provided as a Source Data file.

### Binding of Stat1 to GAS motifs activated *bmp8a* expression

To explore how virus induces expression of *bmp8a*, we searched for the transcription factor binding sites in *bmp8a* promoter region at the web http://jaspar.genereg.net/. Two gamma-activated sites (GAS), 5‘-ATTCCGGGAAA-3’ (P1) and 5‘-TTTACTAGAAC-3’ (P2), in *bmp8a* promoter region were identified (Supplementary Fig. 4). In mammals, the transcription factor STAT1 homodimer was known to bind to the GAS motif in the promoter region of IFN-γ triggered downstream genes^11,16^. We thus wonder if Stat1 can bind to the *bmp8a* promoter region to activate the expression of *bmp8a*. In zebrafish, there exist two Stat1 factors, Stat1a (GenBank accession number NM_131480.1) and Stat1b (GenBank accession number NM_200091.2). Thus we constructed dominant-negative mutant plasmids of stat1a-ΔC and stat1b-ΔC, that were transferred into ZFL cells, respectively. It was found that the expression of *bmp8a* was significantly blocked in both cases (Fig. 8a, b). By contrast, the overexpression of either *stat1a* or *stat1b* markedly promoted *bmp8a* promoter-driven luciferase activities (Fig. 8c). In addition, when one of the GAS motifs in *bmp8a* promoter region was deleted, *bmp8a* promoter-driven luciferase activities were significantly reduced; when both of the GAS motifs were deleted, *bmp8a* promoter-driven luciferase activities were reduced even more significantly (Fig. 8d, e). These suggested that both Stat1a and Stat1b may interact with the GAS motifs.

**Fig. 8.**
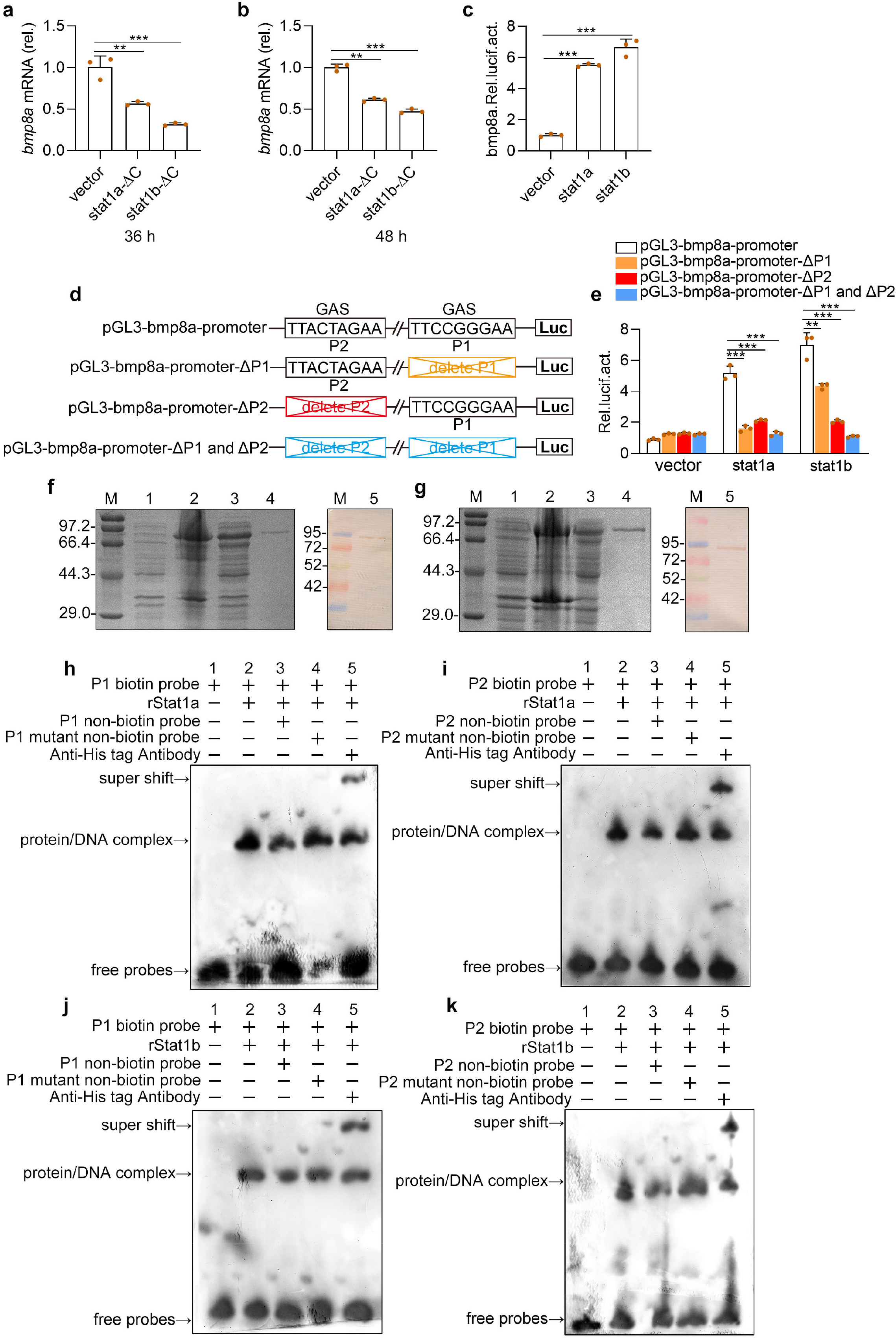
Stat1 binds the GAS sites and activates the *bmp8a* transcriptions upon virus stimulation. **a, b** Expression of *bmp8a* mRNA in ZFL cells after transfected with stat1a-ΔC or stat1b-ΔC for 36 h (**a**) or 48 h (**b**). The expression of *actb1* served as a control for the qRT-PCR. **c** Dual luciferase report assay was used to analyze the transcription abilities of Stat1a and Stat1b in activation of *bmp8a* in EPC cells. pGl3-bmp8a (200 ng) was transfected into EPC cells together with stat1a (200 ng), stat1b (200 ng) or empty vector (200 ng). After 48 h, the transfected cells were collected for luciferase assays. **d** Schematic drawing of wild-type and GAS motif mutation Luc-report plasmids. **e** The 200 ng of pGL3-bmp8a-promoter, pGL3-bmp8a-promoter-ΔP1, pGL3-bmp8a-promoter-ΔP2 or pGL3-bmp8a-promoter-ΔP1 and ΔP2 was transfected into EPC cells along with stat1a (200 ng), stat1b (200 ng) or empty vector (200 ng), respectively. After 48 h, the transfected cells were collected for luciferase assays. *Renilla* luciferase was used as the internal control. **f**, **g** SDS-PAGE and Western-blotting analysis of rStat1a (**f**) and rStat1b (**g**). Lane M: protein molecular standard; Lane 1: negative control for IPTG induced *E*. *coli* (without rStat1a or rStat1b); Lane2: induced rStat1a or rStat1b (the whole cell lysate); Lane 3: induced rStat1a or rStat1b (the supernatant); Lane 4: purified rStat1a or rStat1b; Lane 5: western blot analysis of the sample in Lane 4. **h-k** EMSA was performed to validate the interaction of rStat1a or rStat1b with the GAS motif (P1 or P2) in the bmp8a promoter region. Lane 1: negative group; Lane 2: positive group; Lane 3: an excess unlabeled competitor probe; Lane 4: an excess unlabeled competitor probe containing a mutated runt binding site; Lane 5: Super-shift assays were performed by adding antibody against His tag. Data were from three independent experiments (**a–c, e**) or two independent experiments (**f-k**). Data were analyzed by Student’s *t*-test (two-tailed) and were presented as mean ± SD (\*\**p* < 0.01,\*\*\**p* < 0.001). Source data are provided as a Source Data file.

To verity the binding of Stat1a/Stat1b to the GAS motifs in *bmp8a* promoter region, the recombinant proteins of both Stat1a and Stat1b were expressed and purified (Fig. 8f, g), and used for electrophoretic mobility shift assay (EMSA). As shown in Fig. 8h-k, a remarkable band of protein-DNA complex was observed (lane 2). The protein-DNA complex band was obviously weaker in the presence of an excess unlabeled competitor probe, compared to that of biotin labeled probe group (lane 3), but the intensity of the band showed little change in the presence of GAS mutant probes (lane 4). To detect the specific binding, a supershift experiment with the antibody against His tag was performed. It revealed that His tag antibody caused a specific supershift of slower migrating protein-DNA-antibody complex (Fig. 8h-k, lane 5). There was no protein-DNA complex observed in the negative control group (Fig. 8h-k, lane 1). These demonstrated that both Stat1a and Stat1b could directly bind to the GAS motifs in *bmp8a* promoter region, suggesting that *bmp8a* expression was subjected to the regulation by binding of Stat1a/Stat1b to the GAS motifs.

## Discussion

BMPs, canonical multi-functional growth factors, are suggested to play roles in regulation of immune responses, but it remains controversial over their functions in immunity. We show here for the first time that Bmp8a is a previously unrecognized factor involved in the regulation of antiviral immune responses in zebrafish. Evidences supporting this nature of Bmp8a include: Bmp8a inhibits RNA viruses replication *in vitro* and *in vivo*; Bmp8a promotes the expression of antiviral protein genes including type I IFNs and *mx*; the expression of *bmp8a* is induced by CGRV or poly(I:C), and Bmp8a activates TBK1-IRF3-IFN signaling. Bmp8a is a member of the TGF-β superfamily, which includes more than 30 genes encoding TGFβ, BMPs and activins. TGFβ has been shown to have both pro- and anti-inflammatory effects depending on the context it is acting in the immune system^45^. Like TGFβ, BMP members have also been shown to play different immunoregulatory roles^46^. BMP2, 4 and 7 deficiency or knockdown results in partial loss of the thymic capsule, reduced size of thymus and increased *Helicobacter pylori*-induced inflammation, whereas BMP6 inhibits macrophage growth by inducing cell cycle arrest *in vitro*^27,47–50^. The finding that Bmp8a is a positive regulator in antiviral immune responses provides a new angle for our understanding of the functions of TGF-β superfamily members including BMPs.

On one hand, it is generally regarded that the recognition of virus by PRRs elicits the activation of IRF3 and NF-κB, and induces the production of IFNα/β, that then activate JAK-STAT pathway, eventually leading to the production of ISGs such as cytokines and antiviral proteins. On the other hand, BMPs are known to transmit signals through both Smad-dependent pathways (smad1/5/8 and smad2/3 pathways) and Smad-independent pathways (ERK, JNK, and p38 MAPK pathways). Here we clearly demonstrate that Bmp8a promotes the expression of endogenous type I IFNs independent of Smad signaling pathways. Moreover, it is p38 MAPK pathway, rather than ERK and JNK pathways, that is involved in the antiviral responses induced by Bmp8a. Furthermore, we show that Alk6a participates in Tbk1-Irf3/7-Ifn antiviral signaling via interaction with Bmp8a. In addition, Alk6a was found to be required to induce the phosphorylation of p38 MAPK, which is consistent with the findings in mouse that Alk6 knockdown suppresses p38 MAPK phosphorylation^51^. Previous studies have shown that MAPK signaling pathway is crucial in the phosphorylation of TBK1 and IRF3 upon viral infection^42^. Thus, we probably discover a new pathway of Bmp8a signaling in antiviral immune responses that Bmp8a acts as a positive regulator through the promotion of phosphorylation of Tbk1 via p38 MAPK pathway, i.e., Bmp8a binds to Alk6a, which induces p38 MAPK phosphorylation, and in turn enhances Tbk1 and Irf3 phosphorylation, eventually resulting in increased synthesis of type I IFN (Fig. 9).

**Fig. 9.**
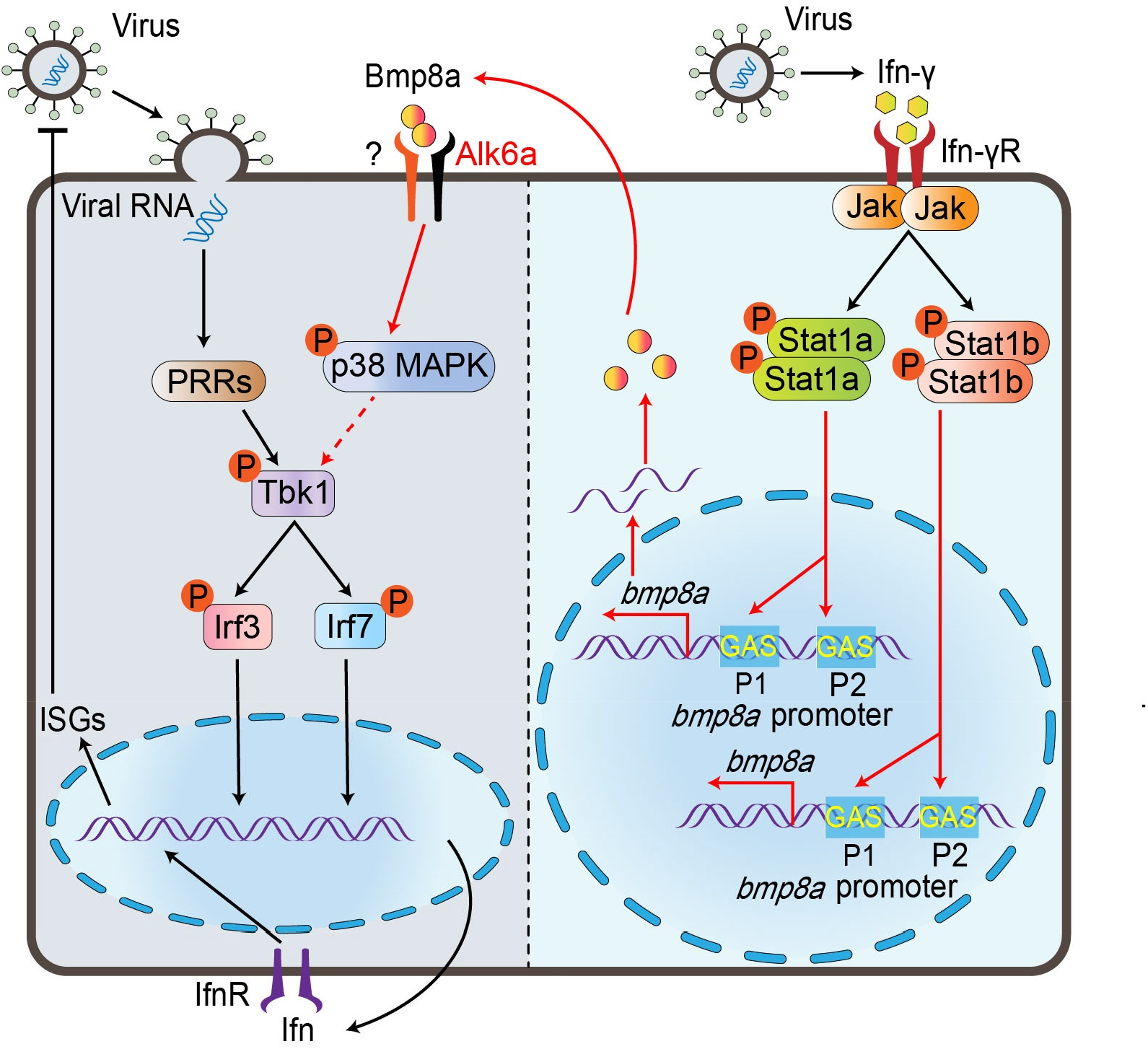
Schematic illustration of Bmp8a serving as positive regulator in the antiviral immune. Upon virus infection, the transcriptions of *bmp8a* are activated through the Jak-Stat1 pathway. The Bmp8a binds to BMP type I receptor Alk6a, promoting phosphorylation of Tbk1 and Irf3 to induce the expression of *Ifn* through p38 MAPK pathway.

Our study also show that the challenge with GCRV and poly(I:C) both significantly promotes the expression of zebrafish *bmp8a*. Searching for the binding sites of transcript factors in *bmp8a* promoter region reveals that two IFN-γ activation sites (GAS) that may interact with STAT1 are identified. Functionally, we demonstrate that Stat1a and Stat1b, orthologues of human STAT1, can directly bind the GAS sites in *bmp8a* promoter, resulting in activation of *bmp8a* expression. It has been reported that in black carp, Stat1a and Stat1b can both form homodimer and heterodimer *in vivo* like their mammalian counterpart^52^. It is thus highly likely that in zebrafish, Stat1a and Stat1b can form homodimer or heterodimer, which then binds to the GAS sites in *bmp8a* promoter region. In zebrafish, IFN-γ has been found to be able to induce the expression of both *stat1a* and *stat1b*, and the GAS motif is required for Ifn-γ activation^53,54^. It is widely known that the JAK-STAT pathway is one of the most important candidate pathways through which IFN-γ works. Activation of JAK-STAT signaling leads to transcription of various downstream ISGs for antiviral activity^55^. We thus think that Bmp8a is another important antiviral factor which can be activated through Ifn-γ-Jak-Stat1 pathway (Fig. 9).

In summary, we demonstrate for the first time that Bmp8a is a positive regulator in antiviral immune responses, which functions through promotion of phosphorylation of Tbk1 via p38 MAPK pathway. Upon virus infection, the expression of *bmp8a* is activated by Stat1a/Stat1b directly. Bmp8a knockdown leads to significantly decreased phosphorylation of Tbk1 and Irf3. As a consequence, *bmp8a*-deficient zebrafish produces significantly lowered type I IFNs in response to RNA virus infection, exhibits significantly reduced immune responses, and increases significantly higher viral load and morbidity *in vivo*. Our study is significant to understand the regulatory mechanism of BMP involving in antiviral immune response and provides a new target for controlling viral infection.

## Methods

### Cells and viruses

The epithelioma papulosum cyprini cells (EPC), zebrafish liver cells (ZFL) were acquired from the China Zebrafish Resource Center (CZRC) and cultured according to the instructions from the CZRC. EPC and ZFL cells were grown in a 28 °C incubator supplied with 5% CO_2_. The flounder gill cells (FG) were grown in a 22 °C incubator supplied. The EPC or FG cells were cultured in MEM media supplemented with 10% fetal bovine serum (FBS) (Gibco), 100 U/ml penicillin and 100 μg/ml streptomycin. The ZFL cells were maintained in DMEM/F-12 media supplemented with 10% FBS, 100 U/ml penicillin and 100 μg/ml streptomycin. Grass carp reovirus (GCRV), a dsRNA virus, and spring viremia of carp virus (SVCV), a negative ssRNA virus, were kindly provided by Yibing Zhang (Institute of Hydrobiology, Chinese Academy of Sciences), and tittered according to the method of Reed and Muench^56^, by a 50% tissue culture infective dose (TCID_50_) assay on EPC cells. The turbot skin verruca disease virus (TSVDV) was isolated from the focal site of turbot skin in our lab.

### Zebrafish

The animals used in the experiment followed the ethical guidelines established by the Institutional Animal Care and Use Committee of the Ocean University of China (permit number, SD2007695). The zebrafish *bmp8a* knockout mutant lines (*bmp8a*^−/−^) were established from the zebrafish AB line using TALENs technology. The TALENs for zebrafish *bmp8a* were assembled using the Unit Assembly (UA) method as previously described^57^. All the UA method starting vectors were the gifts from laboratories of Bo Zhang at Peking University and of Shuo Lin at University of California Los Angeles. TALENs mRNAs were transcribed using the mMESSAGE mMACHINE SP6 kit (Invitrogen, #AM1340) and purifified using the RNeasy Mini kit (QIAGEN, #74106). To generate zebrafish mutant lines, TALEN mRNAs (50–200 pg) were microinjected into 1-cell stage zebrafish embryos. Two days after injection, genomic DNA was isolated from 8 to 10 pooled larvae. The target genomic regions were amplified by nested PCR and subcloned into the pGEM-T vector. Single colonies were genotyped by sequencing. To obtain germline mutations, the TALEN-injected embryos were raised to adulthood and outcrossed with wild-type (WT) fish. The F1 progeny were genotyped by sequencing. To obtain homozygous mutants, heterozygous mutant of the same mutation were obtained and self-crossed. The primers used in this study were listed in Supplemental Table 1.

### Plasmid construction

The open reading frames (ORF) of zebrafish bmp8a, stat1a, stat1b, alk2, alk3, alk6a, actr2a, actr2b, bmpr2a and bmpr2b were cloned into the pcDNA3.1 expression vector for the eukaryotic expression. The ORF of zebrafish stat1a and stat1b were cloned into the pET-28a expression vector for the expression of recombinant proteins. The promoter regions of bmp8a, bmp8a-ΔP1 (delete the GAS motif of 5’-attccgggaaa-3’ in the bmp8a promoter), bmp8a-ΔP2 (delete another GAS motif of 5’-tttactagaac-3’ in the bmp8a promoter), bmp8a-ΔP1and ΔP2 (delete both the two GAS motifs), IFNφ1, IFNφ3, and EPC IFN were cloned into the pGL3-basic vector. Dominant negative mutant plasmids including tbk1-K38M, stat1a-ΔC, stat1b-ΔC, irf3DN, and irf7DN were described previously^58,59^. Dominant negative mutant plasmids alk6a-ΔGS (delete the sequence of 5’-agctccggttctggctcagga-3’ encoding the GS domain) was cloned into the pcDNA3.1 expression vector. The deletions and mutations were created using Mut Express II Fast Mutagenesis Kit V2 (Vazyme, #C214-01). The primers used in this study were listed in Supplemental Table 2.

### Transcriptome Sequencing

The zebrafish were fasted for 24 h prior to tissue collection. Three mutant (*bmp8a*^−/−^) and three wild type (WT) adult zebrafish (0.4 g ± 0.05 g) were randomly selected from each group and were injected intraperitoneally with 50 μl of GCRV (10^8^TCID_50_ per ml) per fish. Seventy-two hours after the injection, the livers from three fish each group were pooled together for total RNA extraction. Three independent samples from each group were prepared for transcriptome sequencing. mRNAs were enriched by magnetic beads combining the oligo (dT). cDNA fragments were subjected to the end repair process by the addition of a single ‘A’ base and the ligation of Illumina sequencing adapters. The ligation products were selected by agarose gel electrophoresis, amplified by PCR, and sequenced using Illumina HiSeq 4000 by Annoroad Technology (Beijing, China).

### Identification of DEGs

The RPKM (reads per kb per million reads) method was used to calculate and normalize the abundances of genes^60^. DEGs across groups were identified by the edgeR package (http://www.r-project.org/). Genes with a fold change ≥ 2 and a false discovery rate (FDR) < 0.05 in a comparison were regarded as significantly expressed. DEGs were then subjected to enrichment analysis of KEGG pathways.

### Viral infection *in vitro*

EPC, FG, or ZFL cells were plated 24 h before infection. Cells were infected with GCRV (5 × 10^4^TCID_50_ per ml) or TSVDV (crude virus extracts filtered by a 0.45 μm microporous membrane) for 1 h in medium without FBS. After infection, cells were washed with PBS and then medium was added with FBS. The cells and cell-free supernatants were harvested at the indicated times.

### Viral infection *in vivo*

For *in vivo* viral infection, the adult WT and *bmp8a*^−/−^ zebrafish were infected via intraperitoneal injection using 50 μL of SVCV (10^8^TCID_50_ per ml), GCRV (10^8^TCID_50_ per ml), or TSVDV (crude virus extracts filtered by a 0.45 μm microporous membrane) per fish. Total RNA was extracted and viral RNA expression in the liver, kidney, intestine, and spleen were examined by qRT-PCR. For the survival experiments, zebrafish were monitored for survival after SVCV, GCRV, or TSVDV infection.

### Crystal violet staining

EPC or ZFL cells were transfected with bmp8a or empty vector plasmid. At 24 h post transfection, cells were treated with GCRV (5 × 10^4^TCID_50_ per ml). At 72 h after virus challenge, cells were fixed with 4% formaldehyde for 1 h and stained with 0.5% crystal violet for 2 h. After washing with tap water, visible plaques were photographed.

### Luciferase assays

EPC cells were seeded in 24-well plates overnight and cotransfected with various plasmids using FuGENE HD Transfection Reagent (Promega, #E2311) following the manufacturer’s instruction. If necessary, the cells were infected with GCRV (5 × 10^4^TCID_50_ per ml). At the indicated time points, luciferase activities were measured with the Luc-Pair Duo-Luciferase Assay Kits 2.0 (iGene Biotechnology, #LF002). Data were normalized by calculating the ratio between firefly luciferase activity and *Renilla* luciferase activity. The primers used for cloning the promoter are shown in Supplemental Table 2.

### RNA interference

Small interfering RNA (siRNA) of bmp8a and si-NC (negative control) were purchased from RiboBio. The oligonucleotide sequences (siRNA) specific targeting *bmp8a* mRNA was as follows: 5’-GTTCTTCAGAGCTAGTCAA-3’. Transfection was performed with Lipofectamine RNAiMAX (Invitrogen, #13778075) according to the manufacturer’s protocols. Total RNA was extracted for qRT-PCR analysis.

### Drug treatment and analysis

SMAD1/5/8 inhibitor DMH1(Selleck, #S7146), smad2/3 inhibitor TP0427736 HCl (Selleck, #S8700), p38 MAPK inhibitor SB203580 (Beyotime, #S1863), JNK inhibitor SP600125 (Beyotime, #S1876), and MEK1/2 inhibitor U0126 (Beyotime, #S1901) were dissolved in dimethyl ulfoxide (DMSO). ZFL or EPC cells were treated with each inhibitor, which were diluted with culture medium at concentrations of 10 μM for 24 h. Total cellular RNA was extracted for qRT-PCR analysis.

### RNA quantification

Total RNA was isolated using the miRNeasy Mini Kit (QIAGEN, #217004) according to the manufacturer’s instructions. RNA treated with DNase and the cDNAs were synthesized with PrimeScript™ RT reagent Kit with gDNA Eraser (TaKaRa, #RP047A). Samples without reverse transcriptase were also added as control. Gene expression was determined by amplifying the cDNA with ChamQ SYBR Color qPCR Master Mix(Vazyme, #Q431-02) by an ABI 7500 Fast Real-Time PCR System (Applied Biosystems). The expression of zebrafish *actb1* or EPC *actin* was used as an internal control. The 2^−ΔΔCt^ method was used to calculate relative expression changes. All qRT-PCR experiments were performed in triplicate and repeated three times. Primers used are listed in Supplementary Table 1.

### Preparation of recombinant protein

The recombinant protein of rStat1a or rStat1b was expressed and purified as previous description^61^. Briefly, the recombinant plasmid (pET-28a-stat1a or pET-28a-stat1b) was transformed into *Trans*B (DE3) chemically competent cell. The positive transformants were incubated in liquid LB medium containing 50 μg/mL kanamycin at 37 °C by shaking at 160 rpm. When the cells grow to OD600 = 0.5, isopropyl β-D-thiogalactoside (IPTG) was added to a final concentration of 1 mM, and incubated at 19 °C with shaking at 120 rpm for 14 h. After incubation, the cells were harvested by centrifugation, re-suspended in PBS, and broken by ultrasound. The supernatant was centrifuged at 12,000 g for 20 min, and the protein was purified by Ni-NTA Sepharose column. The concentration of purified protein was determined by BCA (bicinchonininc acid) method. The purified recombinant proteins were stored at −80 °C for subsequent experiment.

### Electrophoretic mobility shift assay

Electrophoretic mobility shift assay (EMSA) was performed according to the instruction of Chemiluminescent EMSA Kit (Beyotime, #GS009). Oligonucleotides (Supplementary Table 1) were synthesized and biotinylated by Sangon Biotech Company as required and the complementary oligonucleotides were annealed to form the double DNA probe. The recombinant protein was pre-incubated in EMSA/Gel-Shift binding buffer with poly (dI-dC) at 25 °C for 25 min in the presence or absence of Non-Biotin probe (0.5 μM). Then, all samples were mixed with biotinylated probes (0.5 μM) and incubated at 25°C for 30 min. The binding reaction without recombinant protein was set as the negative control. For supershift analysis, anti-His tag antibody was incubated with the reaction mixture for another 30 min after the labeled probe was added. After separation in a native 6% polyacrylamide gel, the free probes and DNA-protein compound were transferred to a Hybond-N^+^ nylon membrane, followed by crosslinking under UV light. After immersion in the blocking buffer, membranes were incubated with streptavidin conjugated HRP for 20 min with shaking and then washed three times with washing buffer. Membranes were exposed briefly in Western Lightning-ECL and then exposure to Automatic X-ray Film Processor (Smpicgg).

### Immunoblot analysis and co-immunoprecipitation

For Immunoblot analysis, cells were lysed in NP-40 buffer (150 mM NaCl, 0.5% EDTA, 50 mM Tris, 1% NP40, proteinase inhibitors, and protein phosphatase inhibitors) for 30 min on ice. For co-immunoprecipitation (co-IP), whole-cell extracts were collected 48 h after transfection and lysed in NP-40 buffer on ice for 30 min. After centrifugation for 15 min at 12,000 × g, 4 °C, supernatants were collected and incubated with Protein A/G PLUS-Agarose (Santa Cruz, #sc-2003) coupled to specific antibodies for overnight with rotation at 4°C. Beads were washed 3 × times with NP-40 buffer. Bound proteins were eluted by boiling 7min with 2 x SDS sample buffer. For immunoblot analysis, immunoprecipitates or whole-cell lysates were separated by 12.5% SDS–PAGE gels, electro-transferred to PVDF membranes and blocked for 4 h with 5% no-fat milk solution, followed by blotting with the appropriate antibodies and detection by enhanced chemiluminiscence (ECL). Antibodies from Cell Signaling Technology: TBK1 (1:1000, #3504T), p-TBK1(Ser172) (1:1000, #5483T). Antibodies from Bioss: IRF3 (1:1000, #bs-2993R), p-IRF3 (Ser386) (1:1000, #bsm-52170R), p38MAPK (1:1000, #bs-0637R), p-p38MAPK (Thr180 + Tyr182) (1:1000, #bs-2210R), Actin (1:2000, #bs-0061R). Antibodies from Beyotime: HA tag (1:1000, #AH158) and Flag tag (1:1000, #AF519). Antibodies from CWBIO: His tag (1:5000, #CW0285) and goat anti-rabbit IgG HRP secondary antibody (1:8000, #CW0103S). Antibodies from Abcam: VeriBlot for IP Detection (1:5000, #ab131366).

### Statistical analysis and reproducibility

Representative experiments have been repeated at least two to three times. All statistical analysis was performed using Graphpad Prism 8.0.1. Data are presented as mean ± SEM. Statistical significance was assessed using the unpaired, two-tailed Student’s *t*-test. Survival analyses were performed using the Kaplan-Meier method and assessed using the log-rank (Mantel-Cox) test. \**p* < 0.05, \*\**p* < 0.01, *** *p*< 0.001; ns, not significant, *p* > 0.05.

## Data availability

All data are available from the corresponding author upon reasonable request and that a Source Data file is available for this article.

## Author contributions

Z.L.and S.-C.Z. conceived and coordinated the project. Z.L., G.J., and S.Z. designed the experiments. S.Z., H.L. and Y.-S.W. performed experiments. H.-Y.L. and Y.W. contributed to establish the *bmp8a* knockout mutant lines (*bmp8a*^−/−^) using TALENs technology. S.Z. performed the statistical analysis. Z.L., S.-C.Z. and S.Z. wrote the manuscript, with input from the other authors. All authors reviewed the manuscript and approved the final version.

## Acknowledgements

This work was supported by the grants of National Natural Science Foundation of China (31572259) and National Key Research and Development Project of the Ministry of Science and Technology(2018YFD0900505).

## References

1. Thaiss, C. A., Levy, M., Itav, S. & Elinav, E. Integration of Innate Immune Signaling. Trends Immunol. 37, 84–101 (2016).

2. Akira, S. & Hemmi, H. Recognition of pathogen-associated molecular patterns by TLR family. Immunol. Lett. 85, 85–95 (2003).

3. Chen, S. N., Zou, P. F. & Nie, P. Retinoic acid-inducible gene I (RIG-I)-like receptors (RLRs) in fish: current knowledge and future perspectives. Immunology 151, 16–25 (2017).

4. Keating, S. E., Baran, M. & Bowie, A. G. Cytosolic DNA sensors regulating type I interferon induction. Trends Immunol. 32, 574–581 (2011).

5. Parvatiyar, K. et al. The helicase DDX41 recognizes the bacterial secondary messengers cyclic di-GMP and cyclic di-AMP to activate a type I interferon immune response. Nat. Immunol. 13, 1155–1161 (2012).

6. Zhou, P. et al. MLL5 suppresses antiviral innate immune response by facilitating STUB1-mediated RIG-I degradation. Nat. Commun. 9, 1243 (2018).

7. Honda, K. et al. IRF-7 is the master regulator of type-I interferon-dependent immune responses. Nature 434, 772–777 (2005).

8. Tamura, T., Yanai, H., Savitsky, D. & Taniguchi, T. The IRF family transcription factors in immunity and oncogenesis. Annu. Rev. Immunol. 26, 535–584 (2008).

9. Li, W., Hofer, M. J., Jung, S. R., Lim, S. L. & Campbell, I. L. IRF7-dependent type I interferon production induces lethal immune-mediated disease in STAT1 knockout mice infected with lymphocytic choriomeningitis virus. J. Virol. 88, 7578–7588 (2014).

10. Schindler, C. & Plumlee, C. Inteferons pen the JAK-STAT pathway. Semin. Cell Dev. Biol. 19, 311–318 (2008).

11. Decker, T., Kovarik, P. & Meinke, A. GAS elements: a few nucleotides with a major impact on cytokine-induced gene expression. J. Interferon Cytokine Res. 17, 121–134 (1997).

12. Fu, X. Y., Kessler, D. S., Veals, S. A., Levy, D. E. & Darnell, J. E. Jr. ISGF3, the transcriptional activator induced by interferon alpha, consists of multiple interacting polypeptide chains. Proc. Natl Acad. Sci. USA 87, 8555–8559 (1990).

13. Levy, D. E., Kessler, D. S., Pine, R. & Darnell, J. E.Jr. Cytoplasmic activation of ISGF3, the positive regulator of interferon-alpha-stimulated transcription, reconstituted in vitro. Genes Dev. 3, 1362–1371 (1989).

14. Levy, D. E., Kessler, D. S., Pine, R., Reich, N. & Darnell, J. E.Jr. Interferon-induced nuclear factors that bind a shared promoter element correlate with positive and negative transcriptional control. Genes Dev. 2, 383–393 (1988).

15. Ren, Y. et al. Deubiquitinase USP2a Sustains Interferons Antiviral Activity by Restricting Ubiquitination of Activated STAT1 in the Nucleus. PLoS Pathog. 12, e1005764 (2016).

16. Shuai, K., Schindler, C., Prezioso, V. R. & Darnell, J. E.Jr. Activation of transcription by IFN-gamma: tyrosine phosphorylation of a 91-kD DNA binding protein. Science 258, 1808–1812 (1992).

17. Bragdon, B., Moseychuk, O., Saldanha, S., King, D., Julian, J. & Nohe, A. Bone morphogenetic proteins: a critical review. Cell Signal 23, 609–620 (2011).

18. Crisan, M. et al. BMP signalling differentially regulates distinct haematopoietic stem cell types. Nat. Commun. 6, 8040 (2015).

19. Wu, M., Chen, G. & Li, Y. P. TGF-beta and BMP signaling in osteoblast, skeletal development, and bone formation, homeostasis and disease. Bone Res. 4, 16009 (2016).

20. Xue, Y. et al. Organizer-derived Bmp2 is required for the formation of a correct Bmp activity gradient during embryonic development. Nat. Commun. 5, 3766 (2014).

21. Yu, Y., Mutlu, A. S., Liu, H. & Wang, M. C. High-throughput screens using photo-highlighting discover BMP signaling in mitochondrial lipid oxidation. Nat. Commun. 8, 865 (2017).

22. Huber, S. & Schramm, C. Role of activin A in the induction of Foxp3+ and Foxp3-CD4+ regulatory T cells. Crit. Rev. Immunol. 31, 53–60 (2011).

23. Martínez, V. G. et al. The canonical BMP signaling pathway is involved in human monocyte-derived dendritic cell maturation. Immunol. Cell Biol. 89, 610–618 (2011).

24. Martínez, V. G. et al. The BMP Pathway Participates in Human Naive CD4+ T Cell Activation and Homeostasis. PloS One 10, e0131453 (2015).

25. Phillips, D. J., de Kretser, D. M. & Hedger, M. P. Activin and related proteins in inflammation: not just interested bystanders. Cytokine Growth Factor Rev. 20, 153–164 (2009).

26. Seeger, P., Musso, T. & Sozzani, S. The TGF-β superfamily in dendritic cell biology. Cytokine Growth Factor Rev. 26, 647–657 (2015).

27. Takabayashi, H. et al. Anti-inflammatory activity of bone morphogenetic protein signaling pathways in stomachs of mice. Gastroenterology 147, 396–406 (2014).

28. Yasmin, N. et al. Identification of bone morphogenetic protein 7 (BMP7) as an instructive factor for human epidermal Langerhans cell differentiation. J. Exp. Med. 210, 2597–2610 (2013).

29. Sivertsen, E. A., Huse, K., Hystad, M. E., Kersten, C., Smeland, E. B. & Myklebust, J. H. Inhibitory effects and target genes of bone morphogenetic protein 6 in Jurkat TAg cells. Eur. J. Immunol. 37, 2937–2948 (2007).

30. Varas, A. et al. Interplay between BMP4 and IL-7 in human intrathymic precursor cells. Cell cycle 8, 4119–4126 (2009).

31. Chen, W. & Ten Dijke, P. Immunoregulation by members of the TGFβ superfamily. Nat. Rev. Immunol. 16, 723–740 (2016).

32. Zhao, G. Q., Deng, K., Labosky, P. A., Liaw, L. & Hogan, B. L. The gene encoding bone morphogenetic protein 8B is required for the initiation and maintenance of spermatogenesis in the mouse. Genes Dev. 10, 1657–1669 (1996).

33. Zhao, G. Q. & Hogan, B. L. Evidence that mouse Bmp8a (Op2) and Bmp8b are duplicated genes that play a role in spermatogenesis and placental development. Mech. Dev. 57, 159–168 (1996).

34. Walton, K. D. et al. Villification in the mouse: Bmp signals control intestinal villus patterning. Development 143, 427–436 (2016).

35. Ying, Y., Qi, X. & Zhao, G. Q. Induction of primordial germ cells from murine epiblasts by synergistic action of BMP4 and BMP8B signaling pathways. Proc. Natl Acad. Sci. USA 98, 7858–7862 (2001).

36. Zhong, S. j. et al. Spatial and temporal expression of *bmp8a* and its role in regulation of lipid metabolism in zebrafish *Danio rerio*. Gene Rep. 10, 33–41 (2018).

37. Robertsen, B. The interferon system of teleost fish. Fish Shellfish Immunol. 20, 172–191 (2006).

38. Zhang, Y. B. et al. The innate immune response to grass carp hemorrhagic virus (GCHV) in cultured Carassius auratus blastulae (CAB) cells. Dev. Comp. Immunol. 31, 232–243 (2007).

39. Derynck, R., Akhurst, R. J. & Balmain, A. TGF-beta signaling in tumor suppression and cancer progression. Nat. Genet. 29, 117–129 (2001).

40. Drummond, A. E. TGFbeta signalling in the development of ovarian function. Cell Tissue Res. 322, 107–115 (2005).

41. Oh, S. P., Yeo, C. Y., Lee, Y., Schrewe, H., Whitman, M. & Li, E. Activin type IIA and IIB receptors mediate Gdf11 signaling in axial vertebral patterning. Genes Dev. 16, 2749–2754 (2002).

42. Shi, Y. et al. Exosomal Interferon-Induced Transmembrane Protein 2 Transmitted to Dendritic Cells Inhibits Interferon Alpha Pathway Activation and Blocks Anti-Hepatitis B Virus Efficacy of Exogenous Interferon Alpha. Hepatology 69, 2396–2413 (2019).

43. Heldin, C. H., Miyazono, K. & ten Dijke, P. TGF-beta signalling from cell membrane to nucleus through SMAD proteins. Nature 390, 465–471 (1997).

44. Shi, Y. & Massagué, J. Mechanisms of TGF-beta signaling from cell membrane to the nucleus. Cell 113, 685–700 (2003).

45. Worthington, J. J., Fenton, T. M., Czajkowska, B. I., Klementowicz, J. E. & Travis, M. A. Regulation of TGFβ in the immune system: an emerging role for integrins and dendritic cells. Immunobiology 217, 1259–1265 (2012).

46. Tu, E., Chia, P. Z. & Chen, W. TGFβ in T cell biology and tumor immunity: Angel or devil? Cytokine Growth Factor Rev. 25, 423–435 (2014).

47. Aleman-Muench, G. R. & Soldevila, G. When versatility matters: activins/inhibins as key regulators of immunity. Immunol. Cell Biol. 90, 137–148 (2012).

48. Bleul, C. C. & Boehm, T. BMP signaling is required for normal thymus development. J. Immunol. 175, 5213–5221 (2005).

49. Gordon, J., Patel, S. R., Mishina, Y. & Manley, N. R. Evidence for an early role for BMP4 signaling in thymus and parathyroid morphogenesis. Dev. Biol. 339, 141–154 (2010).

50. Lee, M. K. et al. TGF-beta activates Erk MAP kinase signalling through direct phosphorylation of ShcA. EMBO J. 26, 3957–3967 (2007).

51. Itoh, S. et al. Trps1 plays a pivotal role downstream of Gdf5 signaling in promoting chondrogenesis and apoptosis of ATDC5 cells. Genes Cells 13, 355–363 (2008).

52. Wu, H., Zhang, Y., Lu, X., Xiao, J., Feng, P. & Feng, H. STAT1a and STAT1b of black carp play important roles in the innate immune defense against GCRV. Fish Shellfish Immunol. 87, 386–394 (2019).

53. Zou, J. & Secombes, C. J. Teleost fish interferons and their role in immunity. Dev. Comp. Immunol. 35, 1376–1387 (2011).

54. Ruan, B. Y. et al. Two type II IFN members, IFN-γ and IFN-γ related (rel), regulate differentially IRF1 and IRF11 in zebrafish. Fish Shellfish Immunol. 65, 103–110 (2017).

55. Boehm, U., Klamp, T., Groot, M. & Howard, J. C. Cellular responses to interferon-gamma. Annu. Rev. Immunol. 15, 749–795 (1997).

56. Reed, L. J. & Muench, H. A Simple Method of Estimating Fifty Percent Endo-Points. Am. J. Hyg. 27, 493–797 (1938).

57. Huang, P., Xiao, A., Tong, X., Zu, Y., Wang, Z. & Zhang, B. TALEN construction via "Unit Assembly" method and targeted genome modifications in zebrafish. Methods 69, 67–75 (2014).

58. Feng, H., Zhang, Y. B., Zhang, Q. M., Li, Z., Zhang, Q. Y. & Gui, J. F. Zebrafish IRF1 regulates IFN antiviral response through binding to IFNϕ1 and IFNϕ3 promoters downstream of MyD88 signaling. J. Immunol. 194, 1225–1238 (2015).

59. Sun, F., Zhang, Y. B., Liu, T. K., Shi, J., Wang, B. & Gui, J. F. Fish MITA serves as a mediator for distinct fish IFN gene activation dependent on IRF3 or IRF7. J. Immunol. 187, 2531–2539 (2011).

60. Mortazavi, A., Williams, B. A., McCue, K., Schaeffer, L. & Wold, B. Mapping and quantifying mammalian transcriptomes by RNA-Seq. Nat. Methods 5, 621–628 (2008).

61. Sun, C., Hu, L., Liu, S., Gao, Z. & Zhang, S. Functional analysis of domain of unknown function (DUF) 1943, DUF1944 and von Willebrand factor type D domain (VWD) in vitellogenin2 in zebrafish. Dev. Comp. Immunol. 41, 469–476 (2013).

62. Qin, L., Wang. Y. G. & Zhang, Z. Multiple verrucous protrusions on skin in turbot Scophthalmus maximus. J. Dalian Ocean Univ. 23, 479–483 (2008) (in Chinese with English abstract).

